# An extinct hummingbird species that never was: a cautionary tale about sampling issues in molecular phylogenetics

**DOI:** 10.1101/149898

**Authors:** Jorge L. Pérez-Emán, Jhoniel Perdigón Ferreira, Natalia Gutiérrez-Pinto, Andrés M. Cuervo, Laura N. Céspedes, Christopher C. Witt, Carlos Daniel Cadena

## Abstract

The Bogota Sunangel (*Heliangelus zusii*) was described based on a historical specimen lacking locality data as a striking–and potentially extinct– new species of hummingbird more than two decades ago. However, it was considered a dubious taxon by some researchers until a molecular study with strong species-level taxon sampling revealed its phylogenetic affinities and validated its status as a distinct species. We reanalysed existing mitochondrial DNA data together with a new data set sampling multiple populations of the Long-tailed Sylph (*Aglaiocercus kingii*), a species broadly distributed in the Andes of South America. In contrast to previous work, we found that *H. zusii* shares a haplotype with specimens of *A. kingii* from the Eastern Cordillera of Colombia, which is phylogenetically nested within a clade formed by populations of *A. kingii* from the Colombian Andes. These results suggest that *H. zusii* is not a distinct species, but is most likely the result of hybridization between a female *A. kingii* and a male of another hummingbird species. These findings highlight the importance of thorough taxonomic and geographic sampling when assessing the likelihood of hybrid origin of an organism, particularly in cases potentially involving wide-ranging species in areas where deep phylogeographic structure is likely.

## INTRODUCTION

The selection of species and individuals for inclusion in molecular analyses critically affects inferences in various fields of systematic biology including phylogenetics [1], phylogeography [2], and species delimitation [3]. Especially in areas such as the Neotropical region where molecular analyses have recovered substantial within-species divergence and unexpected affinities of populations [4], biases resulting from incomplete taxonomic or geographic sampling may importantly compromise the results of analyses aimed at understanding phylogenetic relationships [5]. Here we document surprising results revealing a case in which inferences regarding the validity of a potentially extinct and iconic species of Neotropical bird were likely compromised because within-species variation was not accounted for in phylogenetic analyses evaluating the alternative hypothesis that the only known specimen may represent a hybrid as opposed to a distinct species.

Hummingbirds (Trochilidae) are well known for their propensity to hybridize, with numerous documented records of interspecific hybridization, often involving species in different genera [6, 7]. Hybrid hummingbird specimens are particularly common in natural history collections and have caused substantial taxonomic confusion because they were often described as distinct species by museum-based ornithologists. Therefore, stringent protocols have been established to diagnose hybrid hummingbird specimens and thus avoid treating hybrids as taxa [8]. The application of such protocols has resulted in the diagnosis of numerous hybrids and, consequently, in the consideration of a large number of named species as invalid. **For example, G. R. Graves has** authored no less than 17 papers on hybrid hummingbird diagnoses since he first described his approach [8] until the present [9].

A notable exception to situations in which historical hummingbird specimens were described as distinct species but were later determined to be hybrids is that of an atypical skin purchased in 1909 by Brother Nicéforo María in Bogotá, Colombia, which was later sent to the Philadelphia Academy of Natural Sciences. Upon examining this specimen, which had puzzled ornithologists for decades and lacked precise locality data, Graves [10] concluded that it was not an aberrant individual of a known taxon and ruled out the possibility that it may represent a hybrid. Therefore, he designated the specimen as the holotype of a new species, the Bogota Sunangel (*Heliangelus zusii*), which he described while noting it may well have gone extinct due to habitat destruction, representing a “relic of a lost world” [10]. Despite the careful consideration and rejection of alternative hypotheses for what this specimen might represent [10], its description as a new species was received with skepticism by some researchers [11].

A study analyzing mitochondrial DNA (mtDNA) sequence data for the only known specimen of *H. zusii* largely settled disagreements about its validity as a species [12]. Phylogenetic analyses remarkably indicated that the specimen is not closely related to species of *Heliangelus*; rather, it was found to be included in a clade with two species in the genus *Aglaiocercus* from northern South America (Long-tailed Sylph *A. kingii* and Violet-tailed Sylph *A. coelestis*) and the Gray-bellied Comet (*Taphrolesbia griseiventris*), a species in a monotypic genus endemic to semiarid scrub habitats in north-central Peru [12]. In addition, sequence divergence between the *H. zusii* holotype and specimens of *Aglaiocercus* and *Taphrolesbia* was considered substantial (>5% and 3% p-distance, respectively), validating its status as a distinct species [12]. Accordingly, the South American Classification Committee of the American Ornithological Society presently treats *H. zusii* as valid species [13].

In 2011, news about observations of a striking hummingbird in montane forests of the Reserva Rogitama, located in the Eastern Cordillera of the Andes in departamento Boyacá, Colombia, produced great excitement among ornithologists and birding enthusiasts, who suspected it might correspond to *H. zusii*. However, after careful examination of a single individual that was captured and released, it was concluded that the Rogitama hummingbird was not *H. zusii*; rather, various phenotypic characters suggested that it was a hybrid, with *A. kingii* and Tyrian Metaltail (*Metallura tyrianthina*) hypothesized to be its most likely parents [14]. Given clarity about the identity of the Rogitama bird, *H. zusii* remains a lost taxon with no records other than the type specimen and is considered critically endangered if not already extinct [15, 16].

Intrigued by the finding of the Rogitama bird, we obtained mtDNA sequence data from a feather sample of it to compare it with data from other hummingbird taxa. Upon initial analyses, we were struck to find that the sequence we obtained was remarkably similar to the published sequence of *H. zusii* available in GenBank [12] in the relatively few nucleotide positions in which they overlapped. Considering that phenotypic traits clearly indicate that the Rogitama bird is not *H. zusii*, we began to entertain a new hypothesis, namely that both of these birds are indeed hybrids, with their similar mtDNA indicating a shared maternal species. Our ongoing work on the phylogeography of *A. kingii*, one of the hypothesized parental species of the Rogitama hummingbird [14], allowed us to compare sequences of this species from various regions with those of the Rogitama hummingbird, the holotype of *H. zusii*, and other closely related hummingbird taxa to evaluate this hypothesis.

## MATERIAL AND METHODS

Detailed analyses of the phylogeny and phylogeography of *Aglaiocercus* and near relatives will be published elsewhere. For the purpose of this study, we sequenced the ND2 mitochondrial gene (ND2) for 32 individuals of *A. kingii* and two of *A. coelestis.* Sampling was designed to cover the distribution range of *A. kingii*, but was especially thorough in montane regions of Colombia, considering the geographic origin of the Rogitama hummingbird and, hypothetically, of *H. zusii* (Figure 1a, Appendix A). Details about the laboratory methods are described in the electronic supplementary material. We combined our new data with published sequences of *H. zusii* (GenBank Accession GU166851), *T. griseiventris* (GU166856), *Adelomyia melanogenys* (JF894047), and *Chalcostigma herrani* (EU042536), with the latter two species designated as outgroups [12, 17]. New sequences were deposited in GenBank (accession numbers XXXX-XXXX). Because we were unable to obtain complete ND2 sequences for the Rogitama hummingbird (we obtained 543 base pairs) and the available ND2 sequence of *H. zusii* is also incomplete, overlap in sequence data between these two specimens was restricted only to 71 base pairs. Overlap with sequences of other specimens, however, was much more extensive.

**Figure 1.**
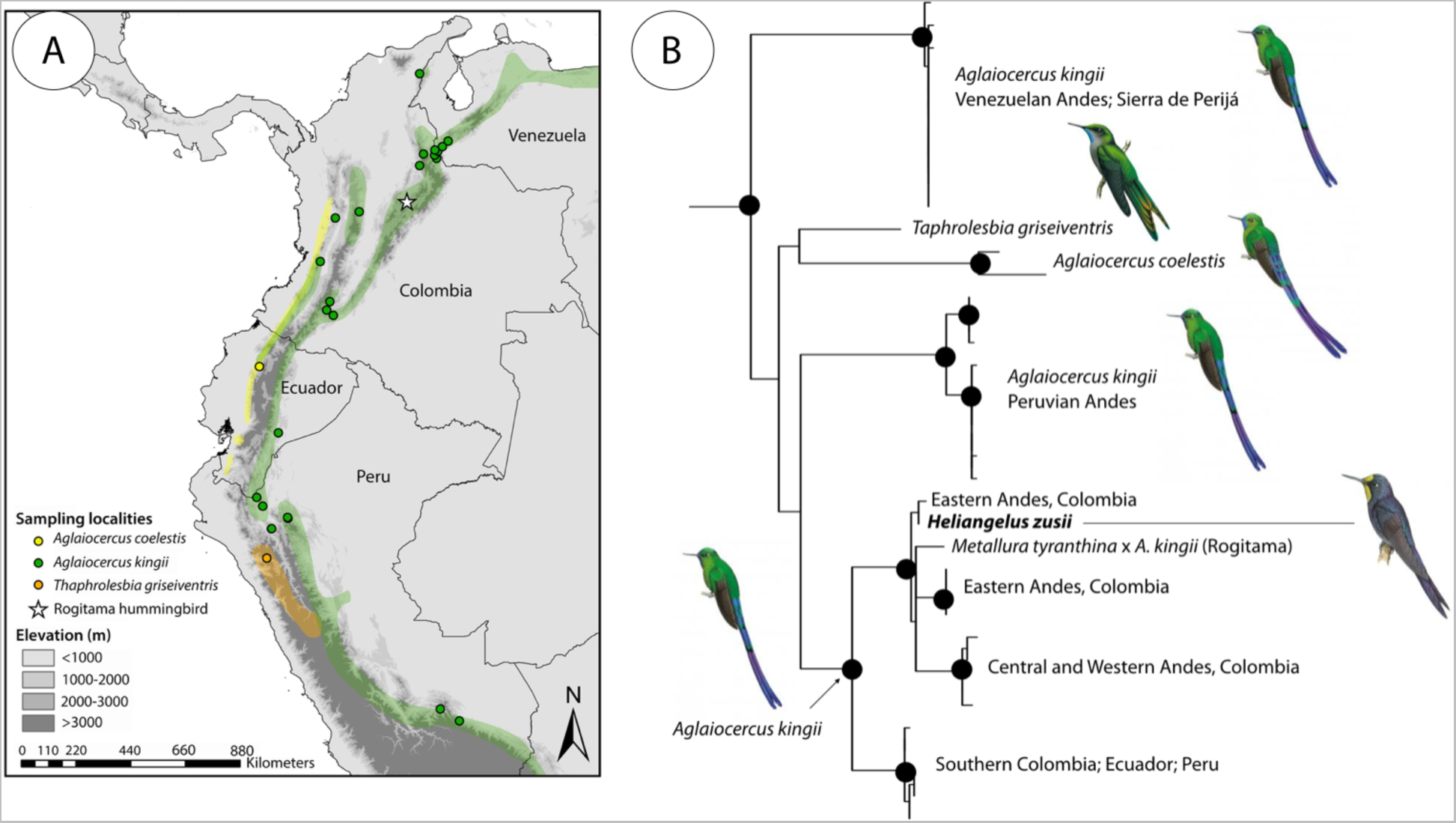
(A) Geographic ranges of *Aglaiocercus kingii*, *A. coelestis*, and *Taphrolesbia griseiventris* in northern South America (polygons), and geographic provenance of specimens of these species and of the Rogitama hybrid hummingbird included in molecular phylogenetic analyses (dots and star). (B) Phylogenetic relationships among species and populations of *Aglaiocercus, Taphrolesbia,* the Rogitama hybrid hummingbird, and *Heliangelus zusii* based on sequences of the ND2 mitochondrial gene. Strongly supported nodes (0.95 Bayesian posterior probability, 80% maximum-likelihood bootstrap) are indicated with black dots. Although nodal support for deep branches is low, note that both the Rogitama bird and *H. zusii* have haplotypes closely allied to those of *A. kingii* from the Eastern Andes of Colombia, indicating they are both hybrids sharing *A. kingii* as female parent. Illustrations courtesy of Lynx Edicions; Handbook of the Birds of the World, Vol. 15, 1999.

We estimated ND2 gene trees using maximum likelihood and Bayesian inference methods. Maximum-likelihood reconstructions were conducted in RAxML-HPC version 8 [18] on XSEDE; we calculated nodal support using 1000 pseudoreplicates using the CIPRES Science Gateway [19]. The Bayesian analysis was carried out using MrBayes v.3.2.6 [20]. We ran four independent runs consisting of four MCMC chains for 10 million generations sampling every 1000 generations, discarding the first 25% of the sampled trees as burn in. We implemented a general-time reversible model of nucleotide substitution with gamma-distributed rate heterogeneity among sites (GTR+G) in both analyses as selected using the Akaike Information Criterion in jModelTest [21].

## RESULTS AND DISCUSSION

Mitochondrial DNA data indicate that *A. kingii* exhibits considerable population structure, with distinct lineages occupying different regions in its distribution range (Figure 1b). We found that sequences of the holotype of *H. zusii* and the Rogitama hummingbird are nested within *Aglaiocercus*, where they clearly belong in a clade formed by individuals of *A. kingii* from the three Cordilleras of the Colombian Andes. Sequences of both “problem” birds are most similar to sequences of individuals from the Eastern Cordillera. In fact, the ND2 sequence of the holotype of *H. zusii* (181 bp) is identical to a sequence of *A. kingii* from Huila (IAvH BT1210; Appendix A) and differs in only one nucleotide from sequences of *A. kingii* from Santander and Norte de Santander (Appendix A). Likewise, the ND2 sequence of the Rogitama hummingbird is identical to sequences observed in individuals from Huila, Santander and Norte de Santander (Colombia), and in individuals from eastern Ecuador and Peru. Indeed, the sequences of *H. zusii* and of the Rogitama hummingbird are identical along the 71 nucleotide sites in which they overlap. Because of the short available sequences of both *H. zusii* and the Rogitama hummingbird, we also analyzed the data considering only the 181 bp fragment we had from *H. zusii* ND2, and results were the same. Phylogenetic analyses of the mtDNA data, however, were unable to recover relationships among clades of *A. kingii* with strong support: major clades formed a polytomy with sequences of *A. coelestis* and *T. griseiventris*, revealing a more complex pattern than suggested by previous phylogenetic studies [12, 17].

We interpret the above the results to imply that, contrary to current views, the holotype of *H. zusii* is probably not a representative of a valid species. Rather, it is most likely a hybrid; because mtDNA is maternally inherited, the data suggest it resulted from a cross between a female *A. kingii* and another species of hummingbird. Given the phylogeographic pattern observed, this hybridization event most likely took place in the Eastern Cordillera of the Colombian Andes. Thus, in addition to resolving the status of an enigmatic specimen, our analysis helps to partly clarify its geographic provenance, considering that “Bogota” trade skins may originate from multiple geographic areas in northern South America [22]. Our results also confirm the hypothesis that the hybridization event producing the Rogitama hummingbird involved a female *A. kingii* [14]. The phenotypic differences between the *H. zusii* and the Rogitama hummingbird suggest that although they share *A. kingii* as female parent they were likely sired by males of different species. However, the observation of phenotypic characters not present in either parental species as shown by recent studies of vocalizations [14] and plumage [23] makes hybrid diagnosis in hummingbirds especially complicated. In fact, our findings underscore the difficulty of diagnosing hummingbird hybrids using phenotypic characters even when rigorous protocols are employed [8, 10]. We thus suggest that analyses of sequences of nuclear genes from the holotype of *H. zusii* would be necessary to establish its other parental species with certainty.

To conclude, we note that our inferred phylogenetic relationships are entirely consistent with those inferred in the study concluding that *H. zusii* is a valid taxon [12], with the radically different conclusion we reached becoming evident only because of our increased taxonomic and geographic sampling. Specifically, because the authors of the earlier study sampled a single individual per species of *Aglaiocercus* (see also [17]), they were unable to detect that *H. zusii* has *Aglaiocercus* mtDNA and to uncover the complex pattern of genealogical relationships among populations of *A. kingii* and with respect to other *Aglaiocercus* and *Taphrolesbia*. Given the unexpected findings of this study and results of other analyses [e.g. 5, 24, 25], we stress that addressing questions about the phylogeny and phylogeography of Neotropical birds–even in cases involving questions about the affinities of a single specimen– requires comprehensive sampling across taxonomy and geography, including multiple individuals per taxon and region.

## Acknowledgements

We thank Roberto Chavarro for permission to obtain feather samples of the Rogitama hummingbird in his reserve and to F. Gary Stiles for providing such samples. We thank the following museums for providing samples for this study: Colección Ornitológica Phelps, Instituto Alexander von Humboldt, Instituto de Ciencias Naturales Colombia, Museo de Zoología of the Universidad Católica del Ecuador, Louisiana State University Museum of Natural Science, and Museum of Southwestern Biology University of New Mexico. Funding was provided by the Consejo de Desarrollo Científico y Humanístico de la Universidad Central de Venezuela (JPE) and University of New Mexico (CW). We thank the national authorities of Colombia and Venezuela for granting research permits. We thank Elisa Bonaccorso for providing sequences of Ecuadorian material and J. Miranda and M. Castro for help with laboratory work. The manuscript was improved thanks to comments by Gary R. Graves.

## Supplementary Material

### Appendix A

Tissue samples sequenced in this study for the mitochondrial ND2 gene. Collection acronyms: Colección Ornitológica Phelps (COP), Instituto Alexander von Humboldt (IAvH), Instituto de Ciencias Naturales Colombia (ICN), Museo de Zoología of the Universidad Católica del Ecuador (QCAZ), Louisiana State University Museum of Natural Science (LSUMNH), Museum of Southwestern Biology University of New Mexico (MSB).

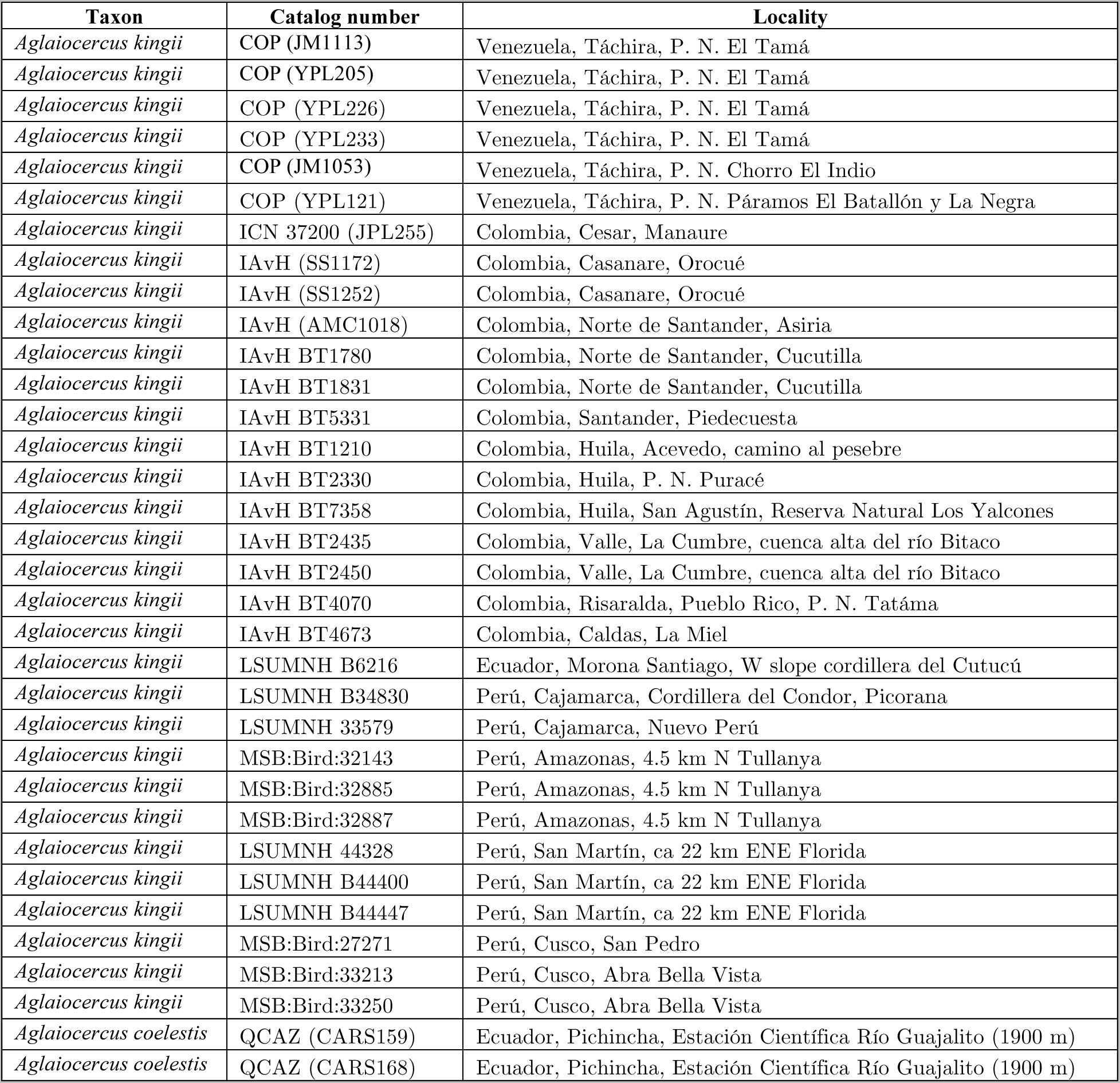

### Laboratory Methods

We isolated whole genomic DNA using the Dneasy Tissue Kit (Qiagen, Valencia, California) following manufacturer’s instructions or a phenol-chloroform method. Amplification of the subunit 2 of the proteing-coding gene NADH (ND2, 1041 bp) was done using the polymerase chain reaction (PCR) in a 2720 termocycler (Applied Biosystems) and primers L5219 and H6313 (Sorenson et al 1999). PCR conditions included an initial denaturation at 95°C for 8 min, followed by 35 cycles of denaturation at 95°C for 30 s, annealing at 50°C for 30 s, and an extension phase of 72°C for 60 s. These cycles were ended with a final extension phase of 72°C for 10 m. The PCR reactions contained 1-2 μL of DNA template, 0.125 U of *Taq* polymerase (PROMEGA), 14.375 μL H20, 1.5 μL MgCl2, 5 μL buffer solution, 1.25 μL of each primer, and 0.5 μL dNTPs, in a total volume of 25 μL. These PCR products were purified in a 1% low-melting point agarose gels and cycle-sequenced using 1 μL DNA template. Molecular work with the Rogitama hummingbird was conducted in a separate lab at Universidad de los Andes and the data for *H. zusii* were obtained from the literature, implying that contamination of samples is not a plausible explanation for our results.

